# Feature Selection via Deep Convolution Neural Networks For Encoding Nonlinear Functional Network Connectivity and Its Application To The Classification of Mental Disorders

**DOI:** 10.1101/2024.09.26.614751

**Authors:** Duc My Vo, Anees Abrol, Zening Fu, Vince D. Calhourn

**Affiliations:** Tri-Institutional Center for Translational Research in Neuroimaging and Data Science (TReNDS), Georgia State, Georgia Tech, Emory University, Atlanta, GA, USA

**Keywords:** schizophrenia, deep convolutional neural network, functional network connectivity, independent component analysis, heatmap

## Abstract

In functional magnetic resonance imaging (fMRI) studies, it is common to evaluate the brain’s functional network connectivity (FNC) which captures the temporal coupling between hemodynamic signals derived from whole brain networks. FNC has been linked to various psychological phenomena. However, analysis of FNCs mainly focuses on linear statistical relationships, which may not capture the full complexity of the interactions among brain intrinsic connectivity networks (ICNs). Therefore, it is important to explore approaches that can better account for possible intricate nonlinear interactions involved in cognitive operations and the changes observed in psychiatric conditions such as schizophrenia. This exploration can lead to a better understanding of brain function and provide new insights into neural links to various psychological and psychiatric conditions. In this paper, we present an innovative approach which utilizes a deep convolutional neural network (DCNN) to extract nonlinear heatmaps from FNC matrices. By analyzing the heatmaps, multi-level nonlinear interactions can be derived from the corresponding input FNC data. Our results show these networks represent a significant improvement over previous approaches and offer a robust framework for understanding the complex interactions between brain regions.

By incorporating two stages in the training process, our method ensures optimal efficiency and effectiveness. In the initial stage, a deep convolutional neural network is trained to create heatmaps from various convolution layers of the network. In the next stage, by utilizing a t-test-based feature selection method, we can effectively analyze heatmaps from different convolution layers. This approach ensures that we are able to functional connectivity with varying degrees of nonlinearity with a focus on the heatmaps that play an important role in distinguishing different groups.

We used a large dataset consisting of both schizophrenia patients and healthy controls, which were divided into separate training and validation sets to evaluate this approach. Results showed patients with increases in default mode networks connections to itself and cognitive control regions and controls with increases within visual and between visual, motor, and auditory domains. We also find significantly increased cross-validated classification accuracy (at 92.8%) compared to several competing approaches. Our approach shows the potential to accurately distinguish differences between the schizophrenia and healthy control groups with high accuracy.

## 1. Introduction

Schizophrenia is a complex and severe mental illness that affects approximately 1% of the global population. It is a chronic and disabling condition that can significantly impact an individual’s quality of life. Patients with schizophrenia often experience a range of symptoms, such as delusions, hallucinations, disorganized thinking and speech, and a lack of motivation and emotional expression. One of the most challenging aspects of schizophrenia is that it can be difficult to diagnose, particularly in its early stages. Some of the early warning signs of schizophrenia can include social withdrawal, reduced speech or difficulty communicating, and a decline in academic or occupational performance. However, these symptoms can also be attributed to other mental health conditions or stressors, making diagnosis a complex and time-consuming process. Early intervention and treatment are crucial for individuals with schizophrenia, as they can help manage symptoms and improve overall functioning. Treatment options may include antipsychotic medications, psychotherapy, and family support. With appropriate care and support, many individuals with schizophrenia are able to lead fulfilling and productive lives.

Functional magnetic resonance imaging (fMRI) has revolutionized our understanding of psychotic disorders such as schizophrenia, providing us with a powerful tool to gain insights into the brain function abnormalities that cause them. Through fMRI, we can now detect and analyze these abnormalities with unprecedented accuracy, helping us to develop more effective treatments and interventions [5]. fMRI studies commonly employ techniques like independent component analysis (ICA) to estimate intrinsic connectivity networks (ICNs) of the brain by analyzing temporal relationships between hemodynamic signals [11] [13]. Researchers have found that understanding the brain’s complex patterns of connectivity and function can provide valuable insights into the underlying mechanisms of cognitive processes. Identifying ICNs is an important part of this research, as ICNs are believed to represent functional sources that play significant roles in various psychological phenomena [8].

Multivariate decomposition techniques, such as ICA, offer a valuable approach to identifying and extracting imaging features. These features include independent components (ICs), time courses (TCs) of independent components, and FNC, which have proven to be highly useful in the study of mental disorders [16] [24]. The TCs offer a valuable representation of the temporal fluctuations in every IC, which is essential for identifying spatially distinct brain regions. Additionally, FNC serves as a crucial tool for characterizing temporal coherence across selected ICs, by correlating their TCs and representing intrinsic connectivity networks. FNC can be used to distinguish individuals with schizophrenia (SZ) from healthy controls (HC) in various predictive clinical contexts, as shown by multiple large-scale meta-analyses. There is a significant body of literature supporting the SZ dysconnection hypothesis. This hypothesis posits that alterations in functional connectivity represent a central endophenotype of the disorder, resulting from various factors including neuromodulatory and synaptic pathogenesis. Most studies of functional connectivity (FC) are designed to estimate networks that reflect linear statistical relationships between different areas of the brain [7] [20] [33]. However, this approach fails to consider the complex nonlinear interactions that are inherent to neural systems. As a result, FC research has largely overlooked the importance of nonlinear interactions in understanding brain function. It is crucial to recognize the significance of non-linear interactions in FC research to gain a comprehensive understanding of the workings of the brain [25]. He et al. [17] demonstrated that the analysis of non-linear statistical relationships is crucial in providing a precise and comprehensive understanding of the organization and dynamics of neural ensembles at multiple scales. By delving into the functional significance of nonlinearity, Friston et al. [16] aimed to unravel the intricacies of information processing in the brain and identify the root causes of mental health disorders. It is because the interplay of neurons in the brain is a crucial factor in shaping our thought processes and actions. Moreover, nonlinear interactions, in particular, can foster a dynamic and versatile environment that facilitates a range of neural computations.

The recognition of nonlinear discriminative patterns and the automatic acquisition of optimal representations from neuroimaging data have made deep learning (DL) methods increasingly attractive for the diagnosis of mental disorders based on fMRI [1] [6] [31]. Due to the ability of DL methods to learn meaningful and complex features from raw data, they have become a promising avenue for enhancing the accuracy and specificity of fMRI-based diagnostic approaches. As such, the application of DL in fMRI-based diagnosis of mental disorders is an area of active research and has the potential to significantly impact the field of psychiatry and neuroscience. Functional connectivity or its network analog, FNC, are widely used input features in deep learning (DL) for analyzing brain function. FC is calculated via the cross correlation among timecourses from a pre-defined brain atlas, whereas FNC is coupling among overlapping whole brain networks, such as those extracted via independent component analysis (ICA). FNC enables the identification of functional relationships between different regions of the brain and can be used to investigate various neurological conditions [37]. Kim et al. [21] conducted a study where they trained a deep neural network (DNN) using functional network connectivity (FNC) data. The DNN was trained with L1-norm to monitor weight sparsity, which refers to the use of a penalty term in the loss function to encourage the model to use fewer features. This approach led to substantial performance improvement in the model’s ability to accurately predict outcomes. Zeng et al. [38] presented a sparse autoencoder to learn imaging site-shared FCs, which was then used to guide SVM training on multi-site datasets for schizophrenia (SZ) diagnosis. To extract and exploit the temporal dynamic information in fMRI time series, researchers have proposed various methods. One such approach is based on recurrent neural networks (RNNs), which are a type of neural network that can process sequences of inputs and maintain an internal state that reflects the context of the previous inputs. By using RNN-based methods, it is possible to model the complex temporal dependencies in fMRI time series and capture the patterns of brain activity over time. These methods have shown promising results in tasks such as brain decoding, brain state classification, and prediction of future brain states. Yan and colleagues [34] [34] have proposed a multi-scale recurrent neural network (RNN) approach to investigate temporal correlations (TCs) in data. On the other hand, Dakka et al. have adopted a recurrent convolutional neural network (R-CNN) on four-dimensional fMRI recordings at the whole-brain voxel level. Their aim was to distinguish patients with SZ and HCs. The proposed approaches have proven to be useful in identifying differences between SZ and HC groups in neuroimaging data. Furthermore, the use of dynamic FNC (dFNC) has become increasingly popular in recent years as a valuable tool in discriminating brain disorders [9] [28] [36]. In some cases, it is used alone, without combining with static FNC, to improve prediction accuracy. This approach involves analyzing the changes in functional connectivity between different regions of the brain over time and can provide valuable insights into how the brain functions and how it is impacted by various disorders. By identifying patterns of connectivity that are unique to specific disorders, researchers hope to develop more effective diagnostic tools and treatment strategies.

The intricate nonlinear interactions exhibited by neural systems are a plausible component of cognitive operations, and it is well-documented that such interactions are altered in schizophrenia. It is imperative to develop highly effective methods for capturing integrated complex networks from measures that exhibit sensitivity towards nonlinear relationships. Our work presents a detailed and innovative approach that utilizes a highly sophisticated deep convolutional neural network (DCNN) [18] to extract nonlinear discriminative heatmaps with remarkable efficiency from the whole training datasets of FNC matrices. Our approach was meticulously designed to overcome the limitations of traditional methods, which often fail to accurately capture the complex nonlinear relationships between different brain regions. By leveraging the power of DCNN, we were able to extract highly informative heatmaps that provide a deeper understanding of the functional connectivity patterns within the brain. Our technique has the potential to revolutionize the field of neuroimaging by providing researchers with a powerful new tool to analyze and interpret complex brain data. This method addresses the challenging task of analyzing brain functional networks using resting fMRI scans, which can vary due to age, sex, lab protocols, and short data acquisition times. Our method involves a two-stage training process. In the first stage, we employ a state-of-the-art deep convolutional neural network to classify FNC training samples into multi-label classes. This process is based on two different kinds of labels, thus allowing for a more comprehensive and practical classification approach. The first label plays a crucial role in accurately classifying and distinguishing between healthy controls and individuals with schizophrenia. The second label is essential in accurately categorizing and differentiating cognitive levels. By maximizing the distances between training FNCs from different classes, we can extract highly nonlinear discriminate features from the heatmaps of DCNN. In the next stage, we plan to implement two highly effective feature extraction methods based on the heatmaps we have generated. These methods have been carefully chosen for their capability to accurately identify and extract key features from the data. The first method involves statistical analysis using t-tests, which are utilized to compare the means of two sets of data. The second method entails the application of ICA to the output of the neural network, with the goal of extracting maximally independent features present in the generated heatmaps. ICA is a computational technique that aims to identify and separate independent sources from a mixture of signals.

## 2. Related Work

Although most studies aim to measure the linear correlation, some researchers have explored nonlinear brain activity modeling. These studies include research conducted by Lahaye et al. [22] in 2003, Stam [29] in 2005, Su et al. in 2013 [30], and Wismüller et al. [32] in 2014. Zwalt et al. [1] showed that, when interpreting fMRI data, it is important to consider the possibility of nonlinearity and its potential impact on the results. It is not common to observe nonlinearity in brain activity, although there are a few known reasons why this might occur. For instance, nonlinear effects of hemodynamic responses in fMRI data can lead to nonlinearity in brain activity. It is worth noting that these effects can vary depending on the time and location of the measurement, as well as between different subjects. Motlaghian et al. [26] [27] indicated that the link between neural activity and large-scale brain networks, as well as the link between networks themselves, is a highly complex and intricate subject of study. The brain is composed of a vast number of individual neurons, which communicate with one another through complex networks of connections. These networks are not static, but rather are dynamic and constantly changing in response to various stimuli. Nonlinearity refers to the fact that the relationship between different brain regions is not linear or predictable, but rather is highly complex and dependent on a multitude of factors. Therefore, further research is needed to fully understand the complex interplay between neural activity and large-scale brain networks, as well as the degree of nonlinearity that exists between these networks. Motlaghian et al. [26], demonstrated that the study of nonlinearity poses several challenges due to the dominance of linear relationships in FNC, which can make it difficult to isolate the effects of nonlinear dependencies. Approaches that focus on the overall relationships may not provide us with a clear picture of the properties of the nonlinear effects. For instance, we may not be able to determine whether these effects are modular, i.e., consisting of independent modules that interact with each other, or manipulable, i.e., capable of being controlled or influenced in a predictable manner. By understanding the modular or manipulable nature of nonlinear effects, we can gain insight into the underlying mechanisms that govern complex systems and develop better models and tools to address real-world problems. Motlaghian et al. [26] tackled this issue head-on by removing the linear relationship and precisely measuring the remaining dependencies, which were the explicit nonlinear dependencies. To accomplish this, they utilized normalized mutual information (NMI). In our case we are focus on the use of nonlinear models to perform predictions and visualizations from precomputed FNCs.

In recent years, there has been a growing interest in studying the dynamics in functional network connectivity (dFNC) over time Allen et al. [4]; Hindriks et al. [19], Lindquist et al. [23]. This line of research aims to gain a deeper understanding of the underlying properties of brain activity and to uncover how different areas of the brain interact with each other in a time-resolved analysis. To explore this phenomenon, researchers have employed an approach where the time courses are divided into smaller windows. By doing so, they can measure the temporal coherence between signals within each successive window. This allows for a more detailed analysis of how the brain functions over time, as opposed to just looking at static snapshots of brain activity. By employing this approach, researchers hope to capture additional insights into the complex nature of the human brain and its functional organization. Ultimately, this research has the potential to shed light on how alterations in brain connectivity may contribute to the development of various neurological disorders, and may ultimately lead to new treatments and interventions.

## 3. ResNet Networks

The objective of our study was to develop a deep learning model for classifying SZ using ResNets [17] - a type of deep residual network. ResNets have revolutionized the field of deep learning by allowing us to create much deeper neural networks, which in turn have led to significant improvements in tasks such as image classification, object detection, and natural language processing. This is achieved by using residual blocks that allow the network to learn the difference between the input and output, thus enabling the network to learn more complex features. Our study utilized ResNets for training the deep learning model for SZ classification, and we observed improved performance in comparison to other models previously used for this task. The use of ResNets in our study thus highlights their potential for enhancing the accuracy and efficiency of deep learning models in various fields. ResNets are a type of deep neural network that address the vanishing gradient problem. This problem arises when training a deep neural network because the gradient signal that is transmitted back through the network during the backpropagation process can become too small to be useful. When this happens, the weights in the network are not updated properly, leading to slower convergence and lower accuracy. ResNets introduce an innovative approach to solving this problem by implementing a ”skip” or ”residual” connection that allows data to bypass network layers. This connection acts as a shortcut that allows the gradient signal to flow more easily through the network, which helps to prevent the vanishing gradient problem from occurring. Specifically, the residual connection allows the network to learn the difference between the input and the output of a set of layers, which can be added back to the output to improve the network’s overall accuracy.

ResNet includes a number of residual blocks (ResBlocks), convolutional layers (Conv), and fully connected layers (FC), as presented in Figure 1. ResBlocks learn residual functions with reference to the layer inputs. This skip-connection approach means that gradients flow more easily through the network, preventing the vanishing gradient problem that can plague deep neural networks.

**Figure 1:**
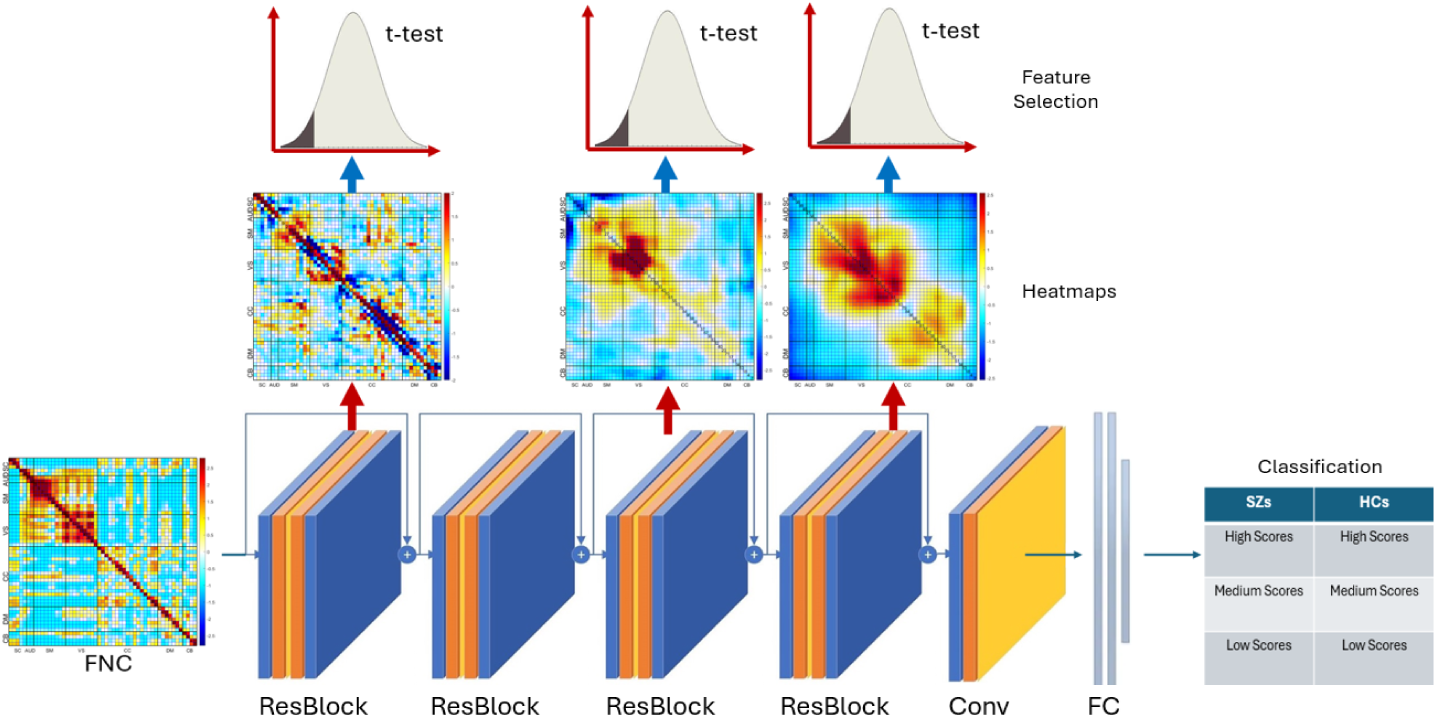
The entire architecture of our proposed method.

In this study, we utilized ResNet as a tool for capturing multi-level features of SZs. ResNet was employed for its ability to extract intricate details that indicate the similarity of SZs to healthy controls (HCs). Through ResNet, we were able to extract high-level features from heatmaps, which contained the most critical connectivities related to SZ. The utilization of ResNet allowed for a comprehensive analysis of the complex features present in SZs, ultimately leading to a more thorough understanding of the disorder and its potential similarities to HCs.

## 4. T-test-based Feature Selection

T-test feature selection is a powerful statistical technique that is used to extract meaningful insights from a dataset. It involves analyzing each feature’s relationship with the target variable and selecting the ones with the strongest correlation. In our analysis, we used a one-sided t-test for selecting features. This test is a type of hypothesis testing that compares the means of two groups and determines whether there is a significant difference between them. In the context of feature selection, it helps to identify which features have a significant impact on the target variable, and which ones can be safely ignored.

The one-tailed t-test is a statistical method that helps to evaluate the significance of the difference between two groups. It is used to test the null hypothesis that there is no statistical difference between the means of two groups against an alternative hypothesis that there is a difference between the means. The critical area of the distribution is one-sided, meaning that it can be either greater than or less than a specific value, but not both.

In our study, we aimed to determine whether the population mean of SZs was significantly lower than that of HCs in a number of intrinsic networks of the brain. We used the one-tailed t-test as it allowed us to identify the most critical function connectivities related to SZ disease in each heatmap. By analyzing the t-value, we were able to determine the statistical significance of the difference between the two groups. Our findings showed that there was a significant difference between the population mean of SZs and HCs in each intrinsic network of the brain, indicating that individuals with schizophrenia have altered functional connectivity in their brains. This information is crucial as it helps us understand the underlying mechanisms of SZ and can be used to develop better treatment methods for individuals with schizophrenia.

## 5. Cognitive Scores

Schizophrenia is a complex mental disorder that can impact an individual’s cognitive abilities.The aim of the study was to explore the cognitive abilities of individuals diagnosed with schizophrenia in comparison to healthy individuals with similar demographic characteristics. Cognitive scores were computed based on neurocognitive domain z-scores derived from computerized neuropsychological tests. We used scores collected via the Computerized Multiphasic Interactive Neurocognitive System (CMINDS) [14] as well as the Measurement and Treatment Research to Improve Cognition in Schizophrenia Consensus Cognitive Battery (MCCB). The study involved 175 patients diagnosed with schizophrenia as well as 169 healthy volunteers. The neurocognitive tests were administered to both groups, and the scores based on the results were analyzed. The findings indicated that the order of the schizophrenia domain profile, based on effect size, was as follows: Speed of Processing, Attention/Vigilance, Working Memory, Verbal Learning, Visual Learning, and Reasoning/Problem Solving. Moreover, the study revealed that women showed higher scores in Attention/Vigilance, Verbal Learning, and Visual Learning. However, men showed higher scores in Reasoning/Problem Solving. This finding is noteworthy as it sheds light on the possibility of sex differences in cognitive abilities. Despite this difference, no significant group by sex interactions were found.

Overall, the study provides a comprehensive understanding of the cognitive abilities of individuals with schizophrenia and the possibility of sex differences in cognitive profiles. The findings can contribute to the development of targeted interventions and therapies aimed at improving the cognitive abilities of individuals diagnosed with schizophrenia. In this study we focus on the use of functional neuroimaging data to predict cognitive scores.

## 6. Proposed Approach

In this section, we provide a comprehensive explanation of our approach to detecting schizophrenia using our algorithm. Our team has developed a novel approach which we outline in the flowchat in Figure 2. This flow chart provides a visual representation of the step-by-step approach that we take to ensure optimal results. Our approach is built on four fundamental steps, each of which plays a critical role in the success of our algorithm.

**Figure 2:**
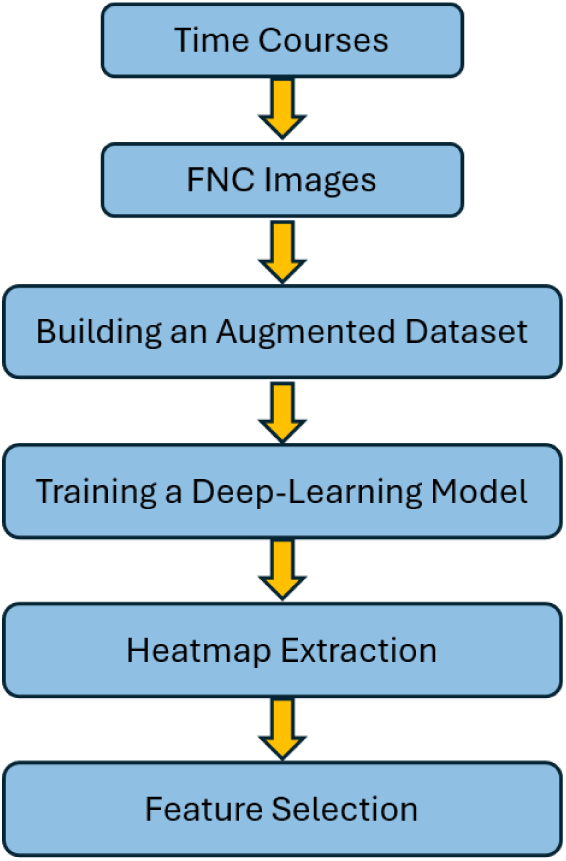
The whole training process of our proposed algorithm.

First, we preprocess the fMRI data and analyze it via an fully automated independent component anlaysis pipeline called NeuroMark [12], resulting in ICNs and their associated timecourses. Each ICN consists of 75 time points. From the timecourses we compute functional network connectivity (FNC), which represent a matrix of the covariance among timecourses.

In the second step of our approach, we focus on presenting various augmentation methods to effectively increase the number of training FNCs based on our initial limited training dataset. These augmentation methods are designed to improve the accuracy and generalization of the FNC models by generating new samples that are similar to the original data but with slight variations. Utilizing an augmented training dataset can be an effective approach towards training a highly efficient residual network. By generating additional synthetic data points, we can increase the size of our training dataset, which in turn can help our model learn more robust and accurate features. These augmentation methods include the mix-up technique. We will discuss this technique in detail and provide examples of how it can be used to enhance the training dataset.

In the third step, the augmented dataset is then used to train a residual network, a type of deep learning architecture that learns to approximate the underlying mapping between the input data and the output labels. Residual networks are known for their ability to train very deep neural networks, which can capture more complex features and patterns in the data. After the training process, we can extract the most powerful and distinguishing deep heatmaps of the model. These heatmaps can be used for various tasks such as classification, detection, or segmentation of SZs. Overall, the use of an augmented training dataset and a residual network can greatly improve the performance and effectiveness of machine learning models in various applications.

In the final step of our approach, we intend to use two highly effective feature extraction methods that are based on the heatmaps we have generated. These methods have been chosen based on their ability to accurately identify and extract key features from the data. The first method of statistical analysis is based on t-tests. T-tests are a type of hypothesis test that is used to compare the means of two groups of data. In this method, the data is first divided into two groups. Then, the mean of each group is calculated, and a t-test is performed to determine whether the difference between the means is statistically significant or not. If it is statistically significant, it means that the difference between the two groups is not due to chance and can be attributed to some other factor. We also conduct a statistical analysis of the intrinsic networks depicted in various types of heatmaps. In order to achieve this, we make use of one-tailed t-tests to compare the means of these networks. This allows us to accurately determine whether the mean of SZ samples in each intrinsic network is significantly greater than or significantly less than the mean of HC samples in every intrinsic network. By performing these tests on each intrinsic network, we can gain a deeper and more nuanced understanding of the complex interrelationships between the different intrinsic networks being studied in the heatmaps. The second method involves the use of a second ICA on the output of the neural network to extract maximally independent features present in the model generated heatmaps. ICA is a computational technique that aims to identify and separate independent sources from a mixture of signals. In the context of heatmaps, ICA is particularly useful as it can help identify the independent sources that contribute to the observed patterns in the data. By decomposing the heatmap into its constituent sources, ICA can help uncover hidden patterns and relationships that may not be apparent from a simple visual inspection of the data.

By employing this rigorous statistical approach, we ensure that our findings are grounded in robust scientific evidence. By following these steps, we will be able to create two highly effective deep learning models that are tailored to two specific objectives. Our first objective is to use a deep convolutional neural network to classify individuals who have been diagnosed with schizophrenia from those who are healthy controls. By doing this, we hope to extract nonlinear discriminative features that can identify and differentiate between the two groups. This DCNN is trained on a large dataset of fMRIs from both SZs and HCs and learns to identify patterns and features that are unique to each group, which can help improve the diagnosis and treatment of schizophrenia. Our second objective is to develop a deep convolutional neural network that can accurately classify each FNC as either a schizophrenia (SZ) patient or a healthy control (HC). This is a significant challenge because FNCs are complex and highly variable. By developing a highly accurate classification model, we can extract discriminative heatmaps that show specific functional connectivity that is predictive of SZ. This may also be useful in tracking changes over the course of the illness and perhaps be useful for developing more effective treatments of interventions. In addition, differentiating between different cognitive levels of SZ and HC can help us understand the relationship between cognition and mental illness along a dimensional scale, using a well defined construct. Overall, our objective is to leverage the power of machine learning to gain new insights into the functional brain patterns linked to SZ.

### 6.1. Preprocessing of Distinct Functional Sources

The human brain is an incredibly complex organ that can be divided into different functional networks or sources, called ICNs. These ICNs interact dynamically with one another to facilitate brain function. ICNs are thought to be responsible for a range of cognitive processes, including memory, attention, decision making, and language processing. Each ICN is made up of multiple regions of the brain that are functionally and anatomically connected. The functional activity of an ICN over time is measured by its time course, which shows how its activity changes over time. The contribution of spatial locations, on the other hand, is indicated by its spatial pattern. To accurately determine the FNC between ICN time courses, we employed Pearson’s correlation coefficient to compute the connectivity between each ICN pair for each individual in our study. This resulted in a 2D symmetric ICN*×*ICN FNC matrix that represented the functional connectivity between two ICNs. The FNC matrix consisted of cells that represented the strength of the functional connectivity between two ICNs. The greater the value of a cell, the stronger the connectivity between the two ICNs. We then use the FNC matrix as input into a deep learning model.

### 6.2. Data Augmentation

The current applicatoin of deep models to study SZ remains a significant challenge due to the lack of sufficient training datasets. This is primarily due to the limitations posed by data accessibility concerns, particularly in light of strict privacy regulations that restrict the sharing of sensitive patient data. As a consequence, insufficient training data significantly affects the performance of SZ classification. In cases where there is an inadequate amount of training data, the accuracy and reliability of the classification model will be significantly reduced, leading to incorrect diagnosis and treatment. Therefore, it is crucial to ensure that an adequate amount of high-quality training data is available to optimize the performance of SZ classification. When the original dataset is limited, it can be challenging to train a machine-learning model. One effective technique to overcome this limitation is data augmentation. Data augmentation refers to the process of generating new training samples by applying various transformations to the original dataset. By generating new samples, data augmentation increases the size and diversity of the training dataset, which leads to better generalization and improved performance of the model. Enhancing the training dataset through data augmentation has been demonstrated to be a highly effective method to reduce over-fitting issues in model training. In this study, we applied mixup augmentation to each ICN. Mixup is a popular and simple data augmentation technique that is widely used in deep learning applications. It involves randomly combining two data points from the same label in the training data set by linearly interpolating between them. This process results in synthetic data points that lie along the straight line connecting the original data points. By generating synthetic data points in the training set, the mixup method helps to increase the diversity of the data and reduce overfitting. One of the key benefits of mixup is that it is easy to implement and does not require any additional labeled data. It has been shown to be effective in a wide range of applications, including image classification, object detection, and natural language processing. The data augmentation steps of our proposed algorithms are presented in Figure 3.

**Figure 3:**
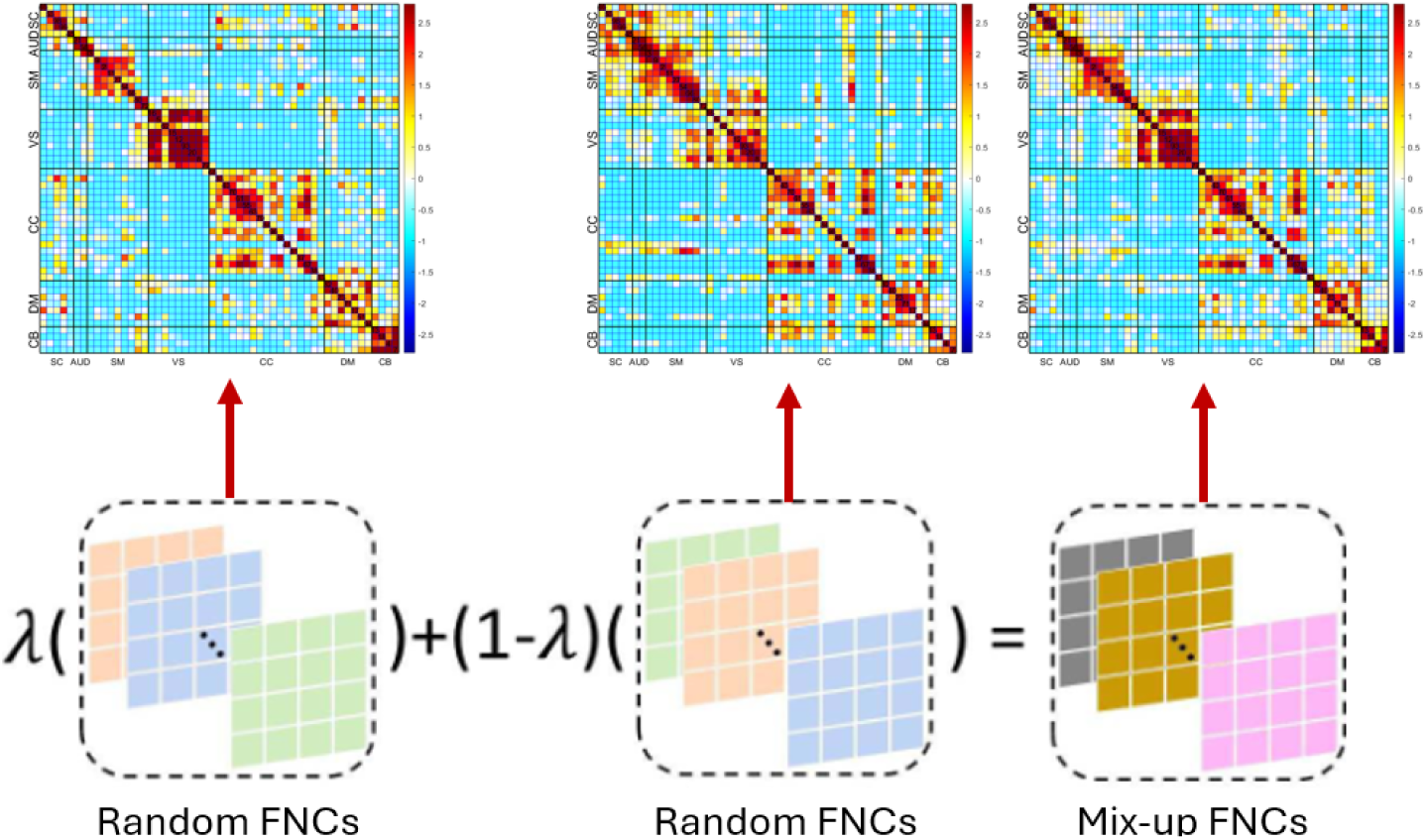
Preprocessing of distinct functional sources.

### 6.3. Training ResNet Models

The ResNet34 network, which was introduced by He et al. [17] in 2016, is used in this study. The network is trained on a dataset that has been augmented to increase the amount of available data. The objective of this study is to classify instances as related or unrelated to schizophrenia. The classification layer is modified to produce a 512-dimensional feature vector, while the feature extraction layer of the ResNet network is kept unchanged. This approach allows for the effective extraction of relevant features from the input data, which can then be used to classify instances accurately. In order to train each network, we make use of the PyTorch distributed machine learning system. To optimize the performance of our model, we utilize a stochastic gradient on a GeForce GTX 1060 Ti. To achieve the highest possible accuracy with minimal error, we set our optimal learning rate to 0.001 and adjust it by decaying every two epochs using an exponential rate of 0.90. This technique allows us to gradually reduce the learning rate, which in turn helps us to converge to the optimal solution. The training process for the ResNet network is conducted over 50 epochs. During this process, the model is trained on the available training data, and its performance is evaluated using a validation set. We make use of various techniques such as data augmentation during the training process to prevent overfitting. By training the model in this manner, we are able to achieve high accuracy and minimize the risk of overfitting. As part of our analysis, we have taken three convolutional layers - layers 10, 26, and 32 - of the model and converted them into heatmaps. These heatmaps allow us to visualize the most critical connections and features that are related to SZ. By providing a detailed two-dimensional score grid, these heatmaps enable us to identify specific regions that play a crucial role in differentiating between SZ and HC classes. This is particularly useful because it allows us to understand how different regions of the brain contribute to the resulting classification. Moreover, by highlighting the regions that are most important for different classes, we can gain a better understanding of the brain regions that differentiate SZ from HC.

### 6.4. Feature Extraction Using T-tests

In order to compare the means of intrinsic networks in heatmaps, we use one-tailed t-tests. One-tailed t-tests are used to determine whether the mean of SZ samples in each intrinsic network is significantly greater or significantly less than the mean of HC samples in every intrinsic network. This statistical method takes into account the variability within each group and calculates a p-value, which indicates the probability of observing the test statistic (t-value) if the null hypothesis is true. If the p-value is less than the predetermined alpha level (usually 0.05), we reject the null hypothesis and accept the alternative hypothesis, which states that there is a significant difference between the means of the two groups. This provides us with a reliable and accurate way to determine whether there is a significant difference between the means of SZ and HC samples in each intrinsic network. In the process of analyzing the neural network’s performance, we have selected specific convolutional layers for further investigation. In order to gain a deeper understanding of the underlying mechanisms at play, we are conducting t-tests on the dataset of heatmaps corresponding to these layers, as seen in Figure 5. The visualization of the mean heatmaps at different convolution layers is presented in Figure 4. The figure is divided into three columns. The first column shows the mean heatmaps of HCs, which are individuals with healthy cognition. The second column presents the mean heatmaps of SZs, which are individuals with schizophrenia. The third column illustrates the difference between the mean heatmaps of HCs and SZs. This difference helps in identifying the regions of the brain where the activity is significantly different between the two groups. The figure has three rows, each depicting mean heatmaps at different layers of convolution. The first row shows examples of mean heatmaps in layer 10, which is a shallow layer. The second row depicts the mean heatmaps in layer 26, which is a mid-level layer. The third row depicts the mean heatmaps in layer 33, which is a deeper layer. The figure provides insight into how the activity patterns in the brain change across different convolution layers.

**Figure 4:**
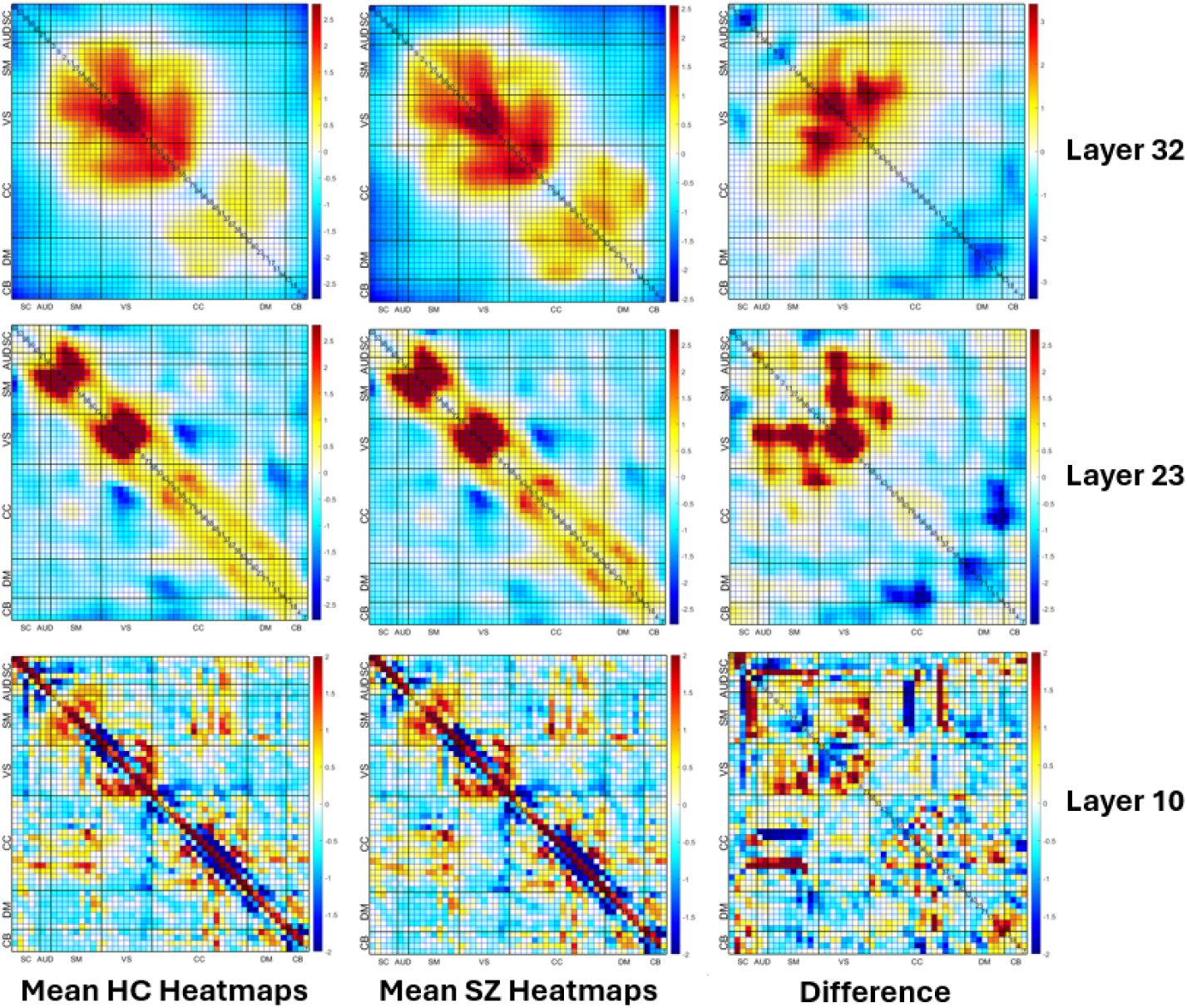
The visualization of the mean heatmaps at different convolution layers. The first column shows the mean heatmaps of HCs, which are individuals with healthy cognition. The second column presents the mean heatmaps of SZs, which are individuals with schizophrenia. The third column illustrates the difference between the mean heatmaps of HCs and SZs. The first row shows examples of mean heatmaps in layer 10, which is a shallow layer. The second row depicts the mean heatmaps in layer 26, which is a mid-level layer. The third row depicts the mean heatmaps in layer 33, which is a deeper layer.

**Figure 5:**
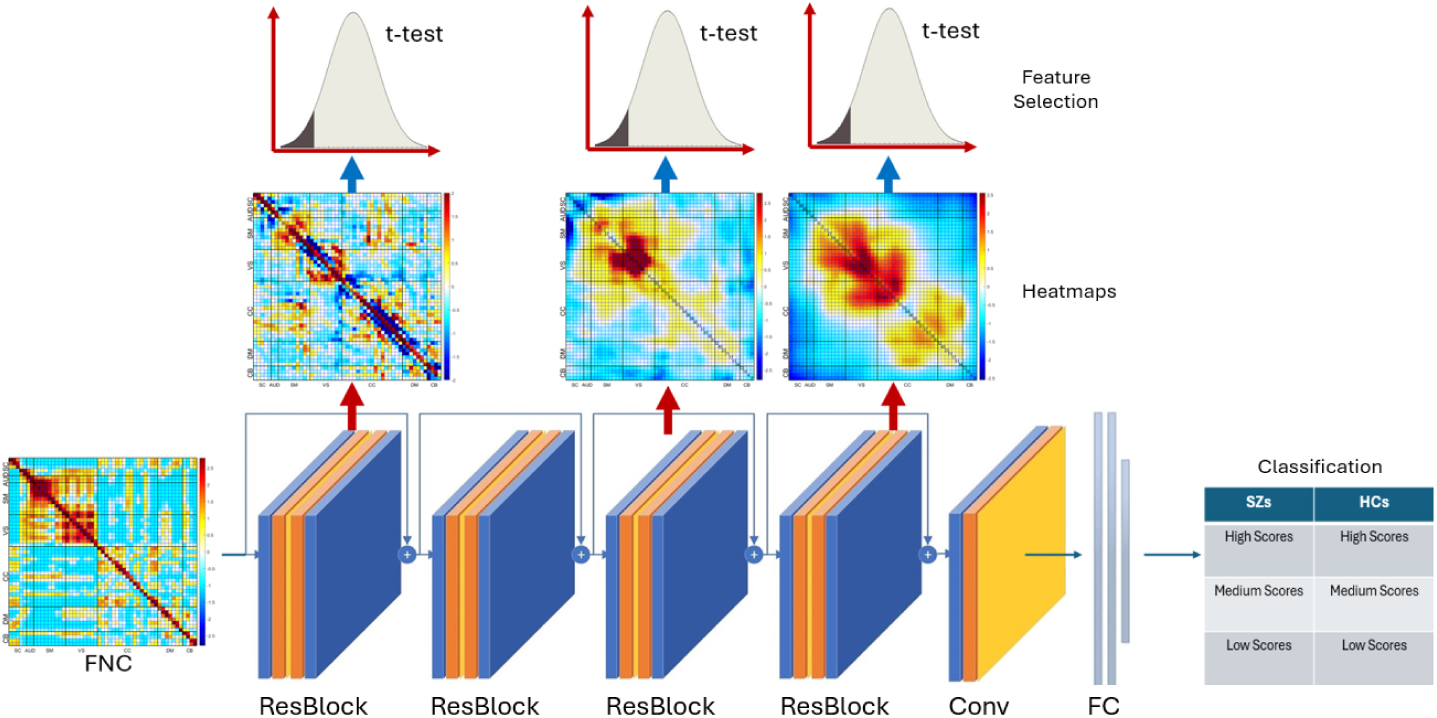
The entire architecture of our feature extraction method using t-tests.

### 6.5. Feature Extraction Using ICA

We take the heatmaps from selected convolutional layers and combining them to generate a comprehensive heatmap that contains all the most prominent deep features. This heatmap helps us better understand the underlying patterns in the data. Figure 6 shows the visualization of combined heatmaps. In this figure, the areas that appear brighter in the heatmaps are indicative of a stronger association with the corresponding class. In this figure, columns 1, 2, and 3 demonstrate the mean heatmaps of SZs, while columns 4, 5, and 6 present the mean heatmaps of HCs. The first row provides a detailed illustration of examples of heatmaps, while the second row depicts the corresponding inputs of the network.

**Figure 6:**
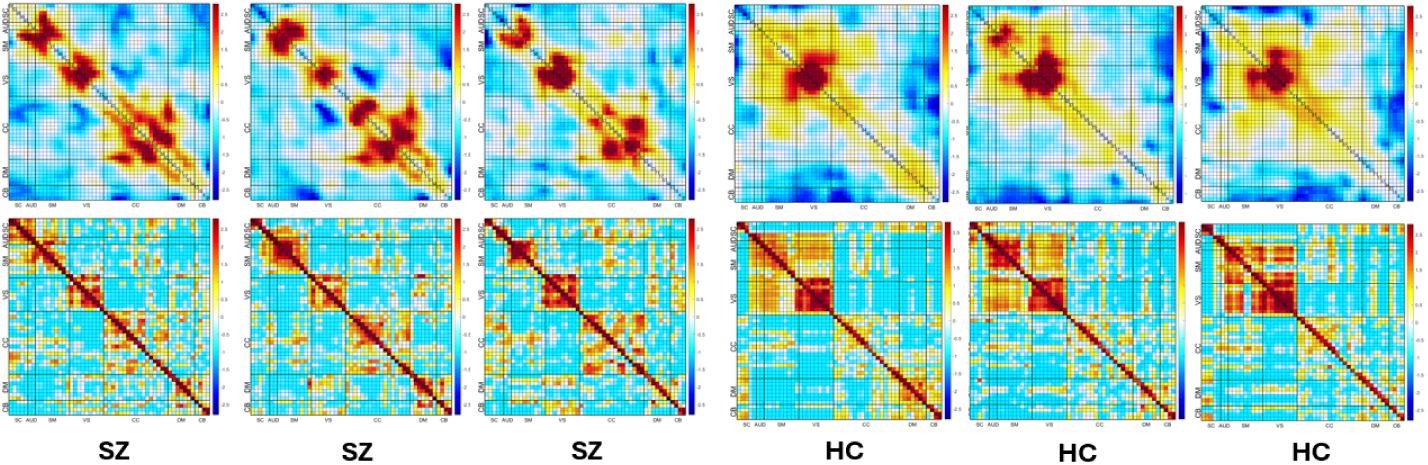
The visualization of combined heatmaps. Columns 1, 2, and 3 demonstrate the mean heatmaps of SZs, while columns 4, 5, and 6 present the mean heatmaps of HCs. The first row provides a detailed illustration of examples of heatmaps. The second row depicts the corresponding inputs of the network.

Next, we apply ICA to this combined heatmap. This is particularly useful in the context of heatmaps because it helps us identify the independent sources that contribute to the observed patterns in the data. This is because ICA can reveal underlying factors that are not directly observable but still contribute to the observed patterns in the data. The combination of heatmaps and ICA allows us to gain a more comprehensive understanding of the deep features in the data and the underlying patterns that they reveal. The architecture of the ICA-based feature extraction is illustrated in detail in Figure 7.

**Figure 7:**
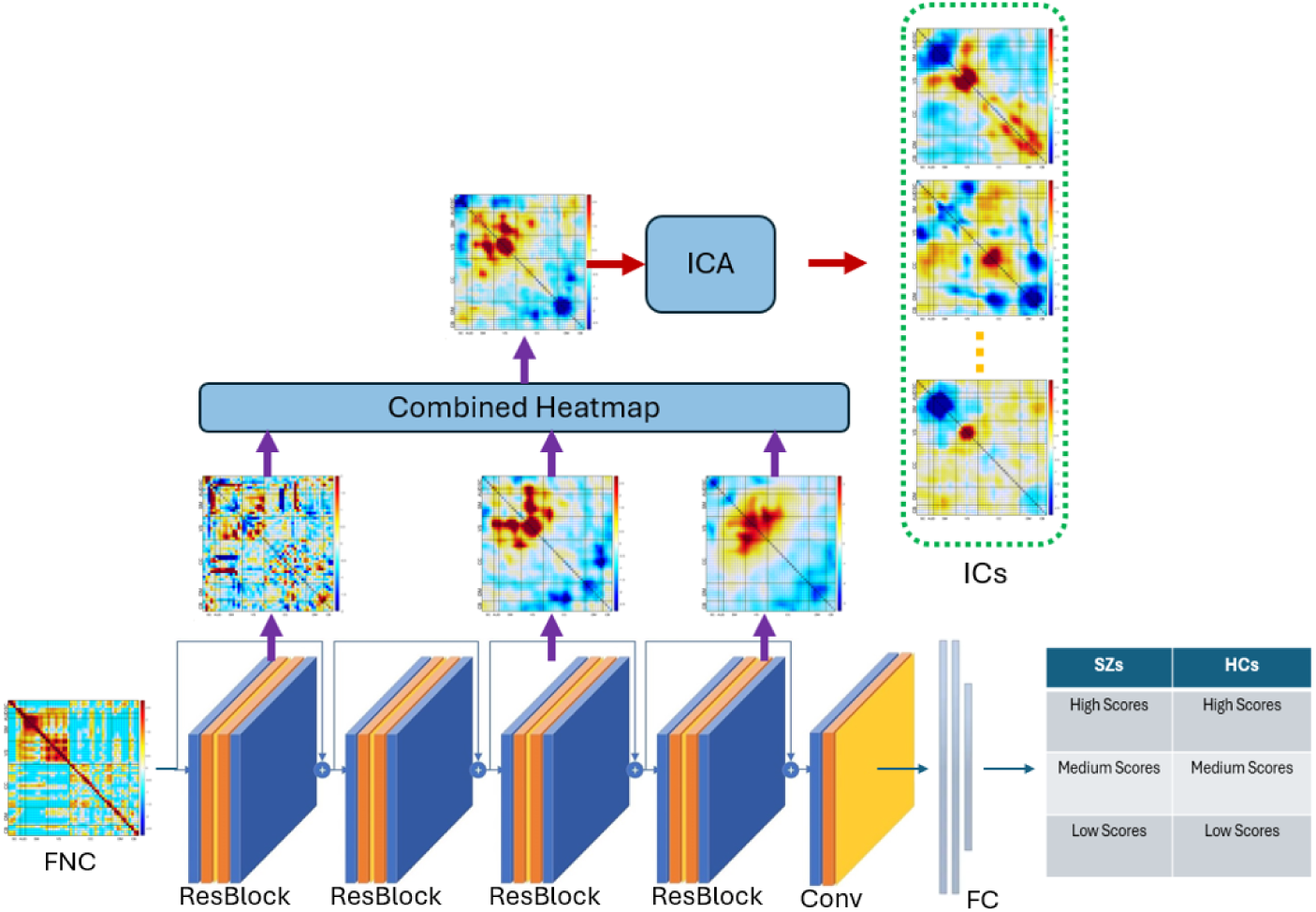
The entire architecture of our feature extraction method using ICA.

## 7. Experimental Results and Analysis

### 7.1. Dataset

In order to evaluate the efficacy of our method, we utilized resting-state fMRI data from a large sample of healthy controls and patients diagnosed with schizophrenia across three datasets - FBIRN (Function Biomedical Informatics Research Network), MPRC (Maryland Psychiatric Research Center), and COBRE (Centers for Biomedical Research Excellence). The dataset included 708 healthy controls and 537 individuals with schizophrenia. Resting-state fMRI data was obtained from the participants while they were in a relaxed state with their eyes closed. The data was then preprocessed and analyzed using advanced statistical and machine learning techniques to identify relevant patterns of brain activity that were predictive of schizophrenia. Recruitment details for the COBRE, FBIRN, and MPRC studies can be located in references Aine et al. [3], Damaraju et al. [10], and Adhikari et al. [2], respectively.

Individuals in the COBRE dataset that had been diagnosed with schizophrenia (SZ) underwent a thorough and rigorous diagnostic process. Two research psychiatrists worked together to make a diagnosis of SZ using the Structured Clinical Interview for DSM-IV Axis I Disorders (SCID), which took into account the patient’s specific circumstances and symptoms. The patient version of the SCID-DSM-IV-TR was used to ensure a standardized approach to the diagnosis. Additionally, the SZ individuals were evaluated for comorbidities that may have been present, and were assessed for both retrospective and prospective clinical stability, to ensure that the diagnosis was accurate and that the patients were receiving appropriate treatment. The participants who were part of the FBIRN study and had SZ (schizophrenia) were diagnosed with the condition using the SCID-DSM-IV-TR (Structured Clinical Interview for DSM-IV-TR). Before undergoing scanning, they were required to be clinically stable for at least two months. For MPRC SZ subjects, a diagnosis of schizophrenia was confirmed via the SCID-DSM-IV.

### 7.2. Experiments

We employed functional magnetic resonance imaging (fMRI) data to extract FNC features for each participant using the NeuroMark independent component analysis pipeline [12]. This pipeline is a fully automated and standardized approach to extract FNC features. Nuisance effects including age, gender, head motion, and site effects were regressed from the data. By using the NeuroMark pipeline and removing nuisance effects from the FNC features, we aimed to ensure that the results we obtained were robust and minimized any potential biases.

To ensure the reliability of our model, we adopted a rigorous approach. Initially, we conducted training and validation using an iterative methodology. The model was trained on the MPRC and COBRE training datasets and subsequently tested on the FBIRN dataset. We expanded the training dataset by adding more augmented training samples and then conducted the test. Following the training phase, we evaluated the model’s performance on the test data.

In order to evaluate how well our two-class classification model performs, we compared it with other classifiers such as support vector machine (SVM), random forests (RF), Kernel SVM (KernelSVM), and decision trees (DT), which are commonly used in the field. To get an accurate assessment of the model’s performance, we used widely-accepted evaluation metrics like precision, recall, accuracy, and F1 score. The results of our analysis are presented in Figure 8.

**Figure 8:**
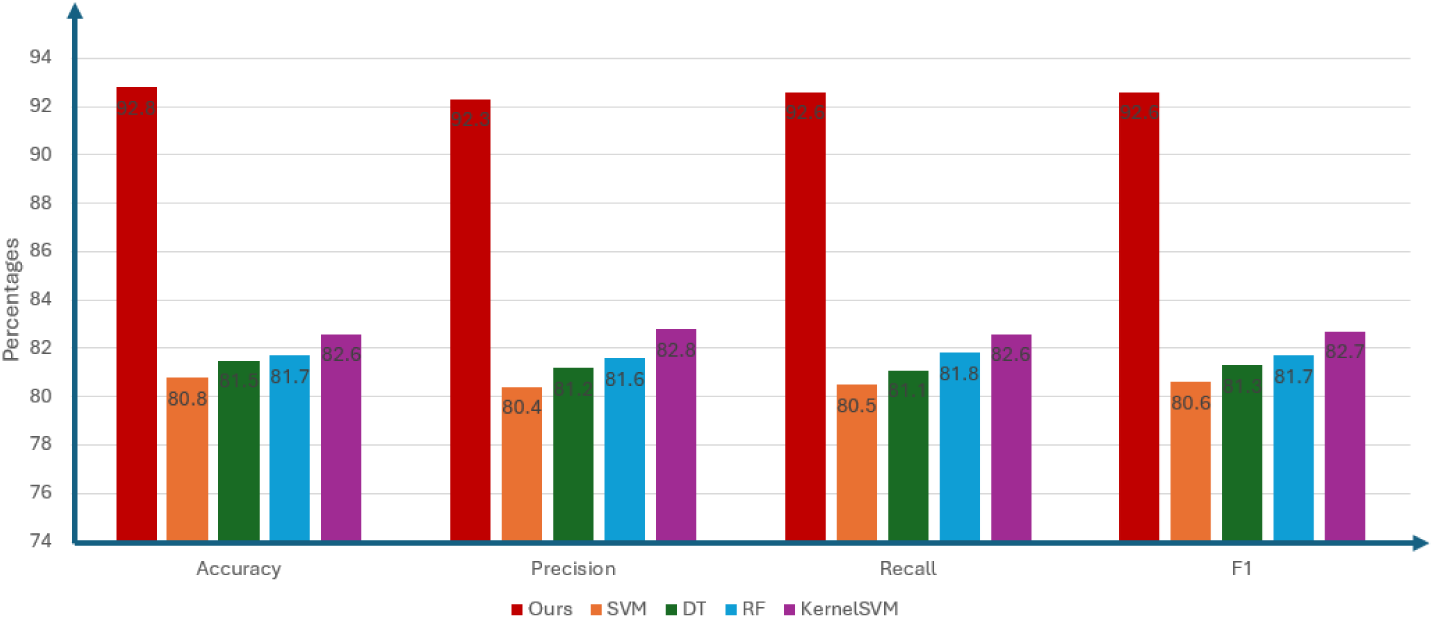
Comparisons of competing methods.

### 7.3. Results

Figure 8 presents our findings, which clearly indicate that our method’s classification accuracy surpasses that of its competitors by a significant margin. The reason for this lies in the highly nonlinear feature space that represents the entire FNC training dataset. As a result, simple linear classifiers like SVM are not effective in classifying these nonlinear features of FNCs. To achieve more precise and reliable results, it is crucial to employ more advanced classification methods that are better suited to the complexity of these nonlinear features.

Our method utilizes ResNet, which is capable of transforming the nonlinear features of FNCs into a learning space that is highly discriminative for classification purposes. This approach enables us to achieve superior classification accuracy compared to other methods. Our approach demonstrated an accuracy rate of 92.8% when classifying patients and controls.

We also utilized an advanced deep convolutional neural network to categorize FNC training samples into multiple classes effectively. This method involved considering two distinct types of labels, which enabled a more thorough and practical approach to classification. The first label was instrumental in precisely differentiating between healthy individuals and those with schizophrenia. The second label was crucial: for accurately classifying and distinguishing cognitive levels. Our approach demonstrated an impressive accuracy rate of 86.5% when applied: to:a multi-class classification task.

### 7.4. Group Differences in Enhanced Heatmaps

The mean heatmaps of healthy controls and individuals with schizophrenia are displayed in Figure 9, with separate columns for each group (the first and second columns, respectively). In addition, the third column shows the difference between the two groups.

**Figure 9:**
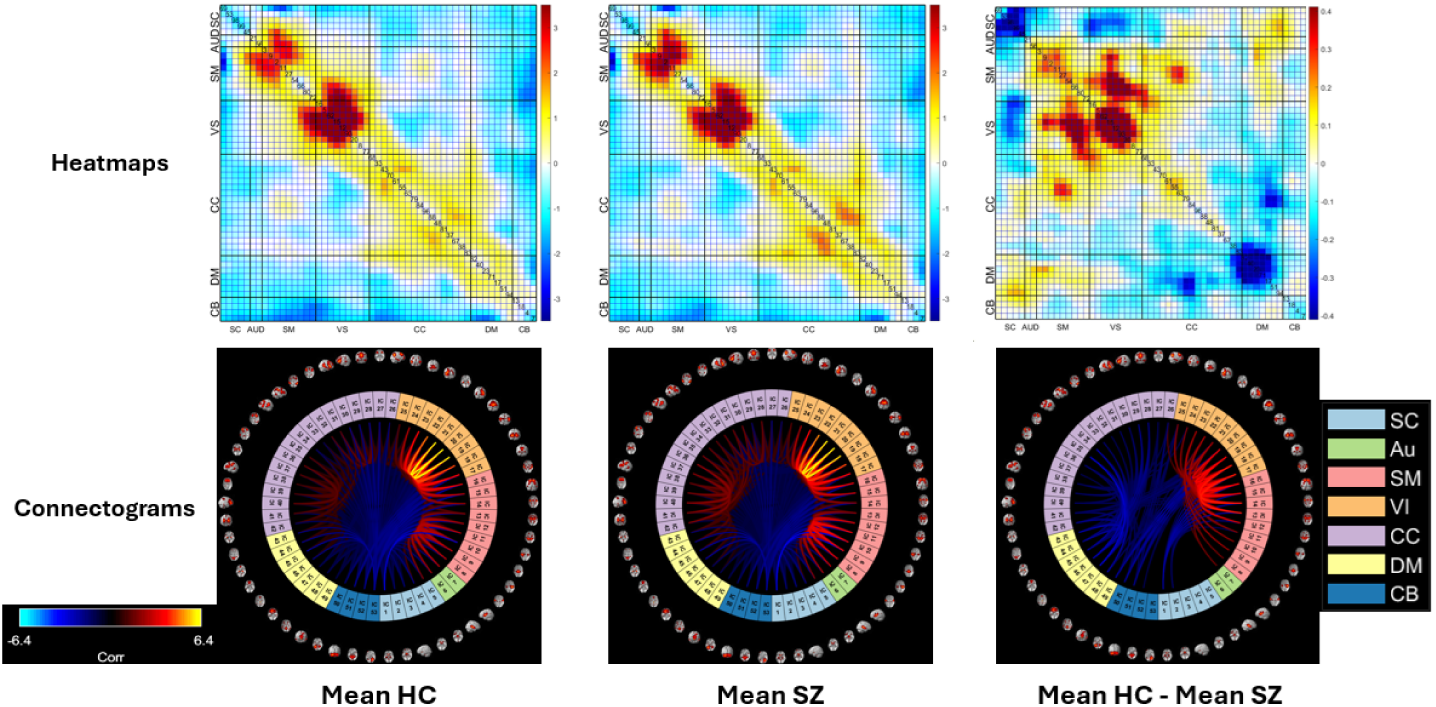
Mean heatmaps of functional network connectivity. The first column shows the healthy controls, the second column shows patients with schizophrenia, and the third column shows the difference between the two groups. The first row shows mean heatmaps, and the second row shows the corresponding connectograms.

We found that the SZ group had significantly weaker connections between three different networks when compared to the HC group. These networks are VS, SM, and CC. This suggests that there may be disrupted neural communication between these regions in individuals with SZ, which could have implications for their cognitive and behavioral functioning. In contrast, the SZ group demonstrated a significant increase in connectivity between the SC network and other networks, such as the AUD, SM, and VS networks, when compared to the control group. Similarly, Figure 9 indicates that there was a marked increase in the level of connectivity between the DM network and the CC network in the SZ group when compared to the control group. This suggests that individuals with schizophrenia may have altered functional connectivity patterns in these particular regions of the brain, which could potentially contribute to the clinical symptoms of the disorder.

The connectograms showcase the most prominent connections present within two distinct networks, VS and SM. The third connectogram, which is located in the third column, shows a significant difference in the strength of connections between the SZ and HC groups. Specifically, the connections in the SZ group are weaker than those observed in the HC group, indicating a disruption in the network connectivity in individuals with schizophrenia.

### 7.5. Group Differences in Different Layers

Our research also aimed to investigate whether there are any significant differences between the means of two distinct population groups: the SZ and HC groups, at different levels of the nonlinear network. Specifically, we conducted two-sample t-tests on layers 10, 22, and 32 to compare the differences between the two groups. By doing so, we can determine whether the differences between the two groups are statistically significant, or if they are just random variations. This analysis helps us to better understand any substantial changes in values and directions between the two groups, and provides additional information regarding the underlying differences between SZ and HC populations.

The analysis presented in Figure 10 comprises the results of two-sided two-sample t-tests conducted for each connectivity within heatmaps at three distinct layers, specifically layers 10, 22, and 32. The null hypothesis for our study states that the mean of the SZ group is equal to the mean of the HC group. We conducted one-tailed two-sample tests to determine whether the mean of the SZ group is significantly greater than the mean of the HC group or if the mean of the SZ group is significantly less than the mean of the HC group, but not both. The first row of the figure displays the heatmaps of two-sample t-tests, which are represented by two different colors. The blue color denotes that the connectivities in the SZ group are weaker than those in the HC group, whereas the red color indicates that the connectivities in the SZ group are stronger than those in the HC group. Additionally, the figure includes the corresponding connectograms that depict the connectivities between regions in a graphical format.

**Figure 10:**
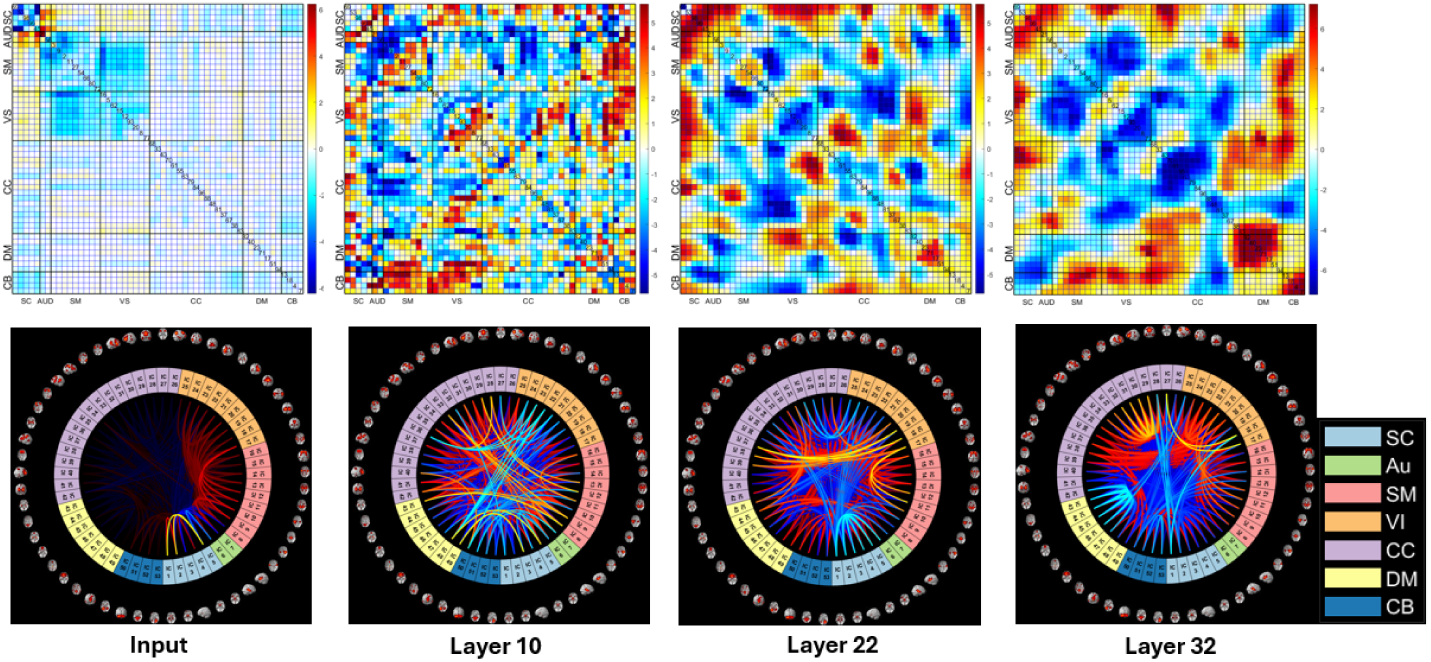
Two-sample t-tests for Inputs, Layer 10, Layer 22, Layer 32. Our research hypothesis assumes that the mean of the SZ group is either significantly greater than or significantly less than the mean of the HC group. The first row shows the heatmaps of two-sample t-tests, which are represented by two different colors. The blue color denotes that the connectivities in the SZ group are weaker than those in the HC group, whereas the red color indicates that the connectivities in the SZ group are stronger than those in the HC group. The second row represents the corresponding connectograms.

The data displayed in this figure indicates that the mean values of the group diagnosed with SZ are significantly lower compared to the group of HC in both between-network connections and within-network connections in the regions of SM, VS, and CC. This implies that there is a notable reduction in the functional connectivity of these regions in individuals with schizophrenia, which could potentially contribute to the manifestation of their symptoms. The SZ group, as compared to the HC group, exhibits stronger connections within the SC network. Additionally, the SZ group also shows evidence of having more robust connections between the SC network and several other networks, such as AUD, SM, VS, and CC networks. In comparison to the HC group, the SZ group also displays significantly stronger connections between the DM network and other networks, including the CC, VS, SM, and AUD networks. These findings suggest that there are distinct differences in brain connectivity patterns between individuals with schizophrenia and healthy individuals. The connectograms highlight the most prominent changes in the connections within three distinct networks: VS, CC, and SM networks. These changes are indicative of the fact that the mean values of the connections in the SZ group are significantly lower compared to the HC group.

The comparison between various layers of ResNet architecture is shown in Figure 10. As per the analysis, it is observed that deep layers, such as layers 22 and 32, demonstrate more discriminative and nonlinear features. In contrast, the input and shallow layer (layer 10) include more linear features along with noise. This behavior is due to the ability of ResNet to eliminate redundant features and noise from the input and retain the most relevant features associated with the classification of SZs and HCs. The deep layers exhibit a high level of abstraction, which helps in capturing complex relationships between different features.

### 7.6. One-tailed Two-sample T-tests on Basis Functional Network Connectivity Components

To further visualize the heatmaps at different nonlinear layers, we applied ICA to the heatmaps from the deep network. Our primary objective was to summarize the results while also capturing the complex nonlinear information via maximally independent FNC maps. This was achieved by identifying a linear transformation of the observed FNCs that maximizes the statistical independence of the resulting components. The process enabled us to apply one-tailed two-sample t-tests on the independent FNCs, which yielded two significant benefits. This allows us to identify which maximally independent FNC regions were most relevant for distinguishing between SZs and HCs. This provides additional insights into the possible underlying neural mechanisms that differentiate SZs from HCs. Second, we could eliminate the connectivities and noise that were irrelevant. This process also helps reduce noise that is less relevant to the SZ classification. This helped us to refine our analysis and improve its accuracy.

In particular, we utilized an ICA model that produced eight basis functional network connectivity components. To determine the statistical significance of the differences between two groups of individuals - those with schizophrenia and healthy controls - we employed one-tailed two-sample t-tests on each of the eight basis FNCs. The null hypothesis for these tests was that the means of independent vectors from the two groups are equal. By performing these statistical tests, we aimed to gain insights into the differences, if any, between the functional connectivity patterns of SZs and HCs. We evaluated the t-tests based on p-values, as shown in Figure 11. Among the t-tests we conducted, the test for the basis FNC component 2 rejected the null hypothesis. This result indicated that the mean of the SZ group was significantly lower than that of the HC group for this particular component.

**Figure 11:**
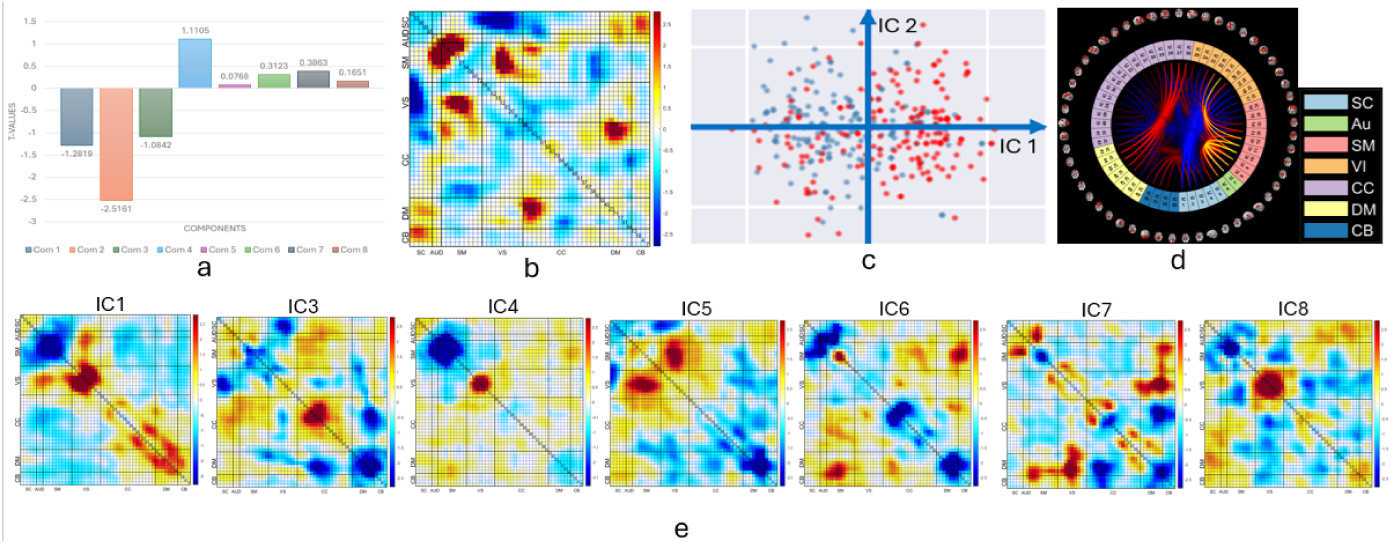
Two-tailed two-sample t-tests on basis functional network connectivity which represents an unmixing of the heatmaps from three network layers. The null hypothesis posits that the mean of the SZ group is equivalent to the mean of the HC group. This one-tailed two-sample test can ascertain whether the mean of the SZ group significantly exceeds the mean of the HC group or if the mean of the SZ group is significantly less than the mean of the HC group, but not both simultaneously. a: P-values for each basis FNC component; b: The second basis FNC component (IC2); c: The projection of the SZ (blue) and HC (red) samples on two axes IC1 and IC2; d: The connectogram of the second basis FNC component; e: This figure displays all the basis FNC components except for IC2.

In Figure 11 b, we can see a heatmap whose loading parameters highlights the FNC values that have the most significant impact on the ability to classify healthy controls and individuals with schizophrenia. This two-tailed two-sample test will determine whether the mean of the SZ group is significantly greater than the mean of the HC group or if the mean of the SZ group is significantly less than the mean of the HC group The FNC cells that exhibit the strongest positive differences (HC-SZ) are the SM and VS networks, as well as the DM and CC networks, as seen in Figure 11 d. These networks have been found to be highly interconnected and play a critical role in regulating various cognitive and affective processes, such as attention, self-awareness, decision-making, and emotional regulation. Additionally, this figure also shows the existence of strong negative connectivitys between the SC and VS networks, which may reflect the inhibitory influence of the SC network on visual processing.

### 7.7. Differences Between The Mean Heatmaps of HCs and SZs At The Same Cognitive Levels

In Figure 12, you can find the advanced heatmaps of both healthy individuals and those with schizophrenia at different cognitive levels. The heatmaps are divided into three groups: high cognitive level (scores ranging from 0 to 2.5), medium cognitive level (scores ranging from -2 to 0) and low cognitive level (scores ranging from -5 to -2).

**Figure 12:**
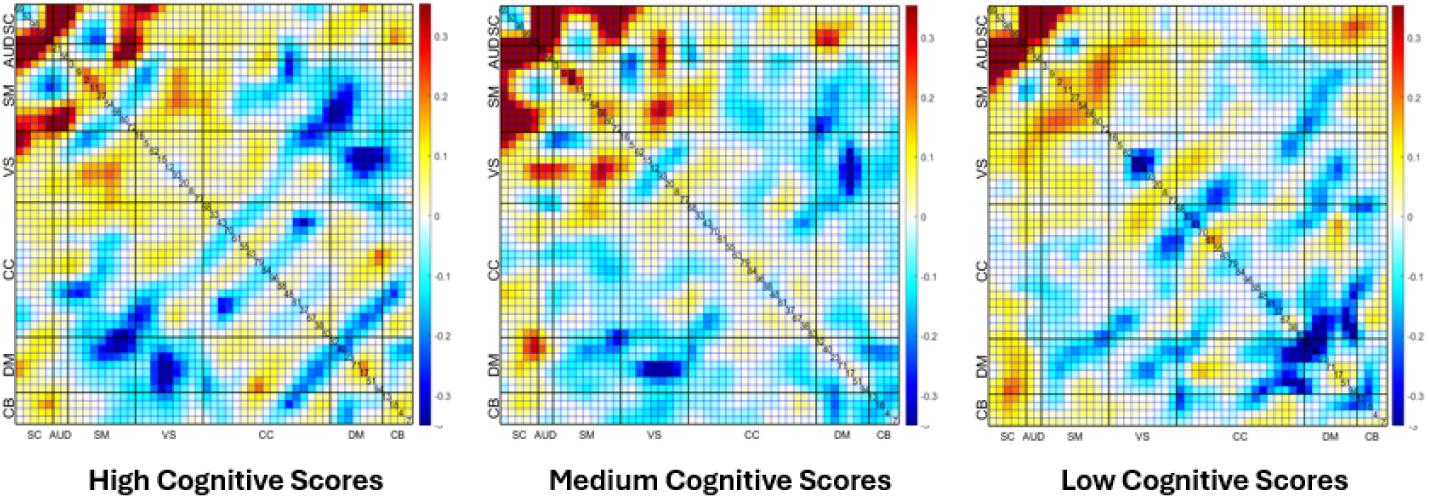
Differences between the mean heatmaps of HCs and SZs at the same cognitive levels.

The results showed that the SZ group had significantly weaker connections between four different networks - the VS, AUD, SM, and SC networks - when compared to the HC group. Further analysis revealed a marked increase in the level of connectivity between the DM network and the CC and VS networks in the SZ group when compared to the control group. This suggests that individuals with schizophrenia may have altered functional connectivity patterns in these particular regions of the brain, which could potentially contribute to the clinical symptoms of the disorder.

### 7.8. Two-sample T-tests on Functional Network Connectivity Components At The Cognitive Levels

In our study, we sought to explore whether there were any significant differences in means between two distinct population groups, SZ and HC, across various cognitive levels. To achieve this, we conducted two-sample t-tests on three different layers, namely 10, 22, and 32, to compare the differences between the two groups at high, medium, and low cognitive score levels. This approach allowed us to determine whether the differences between the two groups were statistically significant or merely random variations. The analysis also enabled us to better understand any substantial changes in values and directions between the two groups at each cognitive level and to draw more reliable conclusions regarding the underlying differences between SZ and HC populations.

The analysis that has been presented in Figure 13 is a comprehensive examination of the results of two-sample t-tests that were conducted for each connectivity found within every heatmap of three distinct layers. These layers include layers 10, 22, and 32, and the t-tests were conducted across three different cognitive score levels, which are represented in three columns respectively. To be more specific, there are three rows in the heatmap, which correspond to the aforementioned layers, while the columns represent the different cognitive score levels that were tested. The heatmaps of two-sample t-tests are represented by two different colors. The blue color denotes that the connectivities in the SZ group are weaker than those in the HC group, while the red color indicates that the connectivities in the SZ group are stronger than those in the HC group. Moreover, the corresponding connectograms that depict the connectivities between regions are shown in Figure 14.

**Figure 13:**
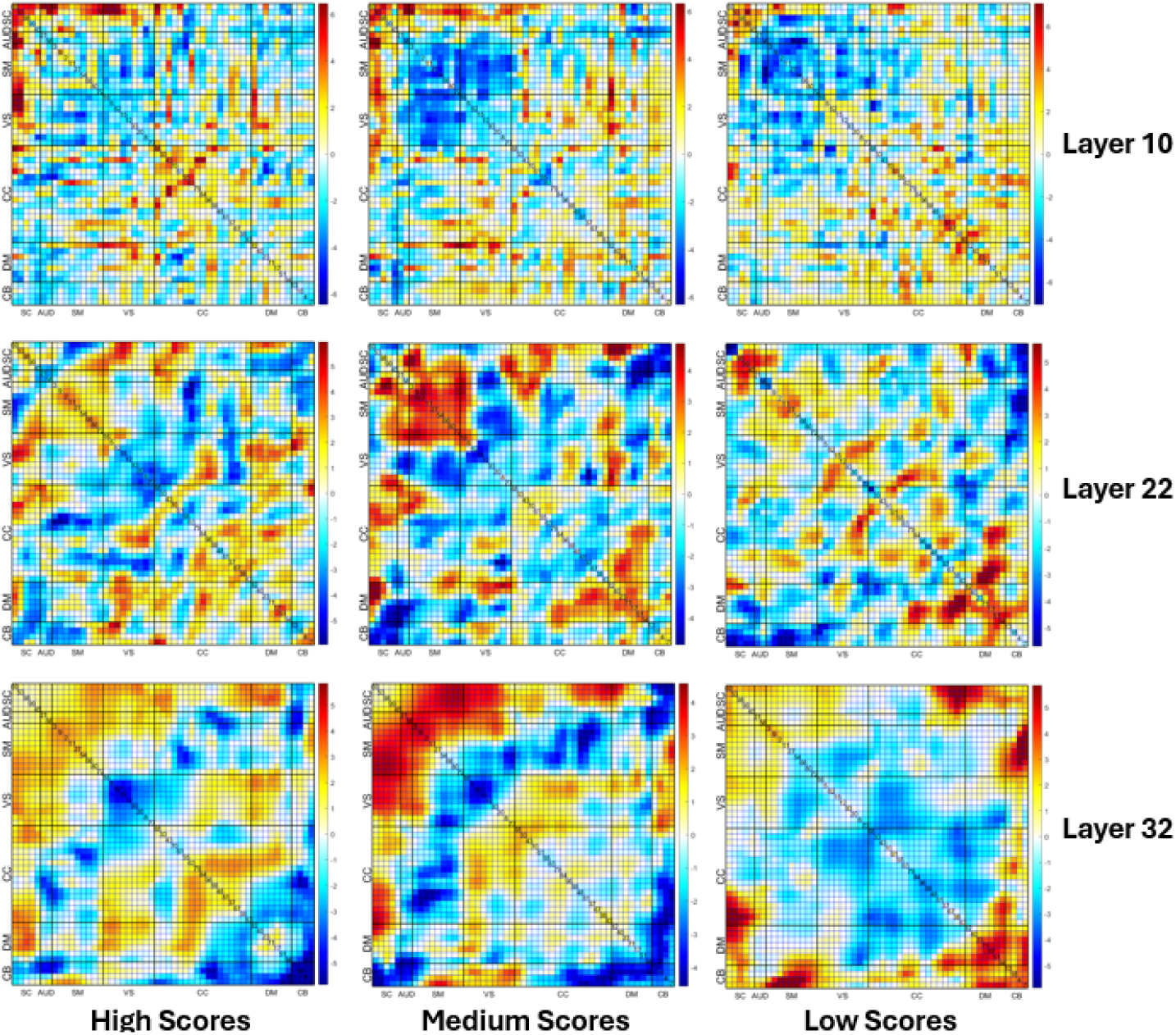
Two-sample t-tests for Layer 10, Layer 22, Layer 32 at different cognitive levels.

**Figure 14:**
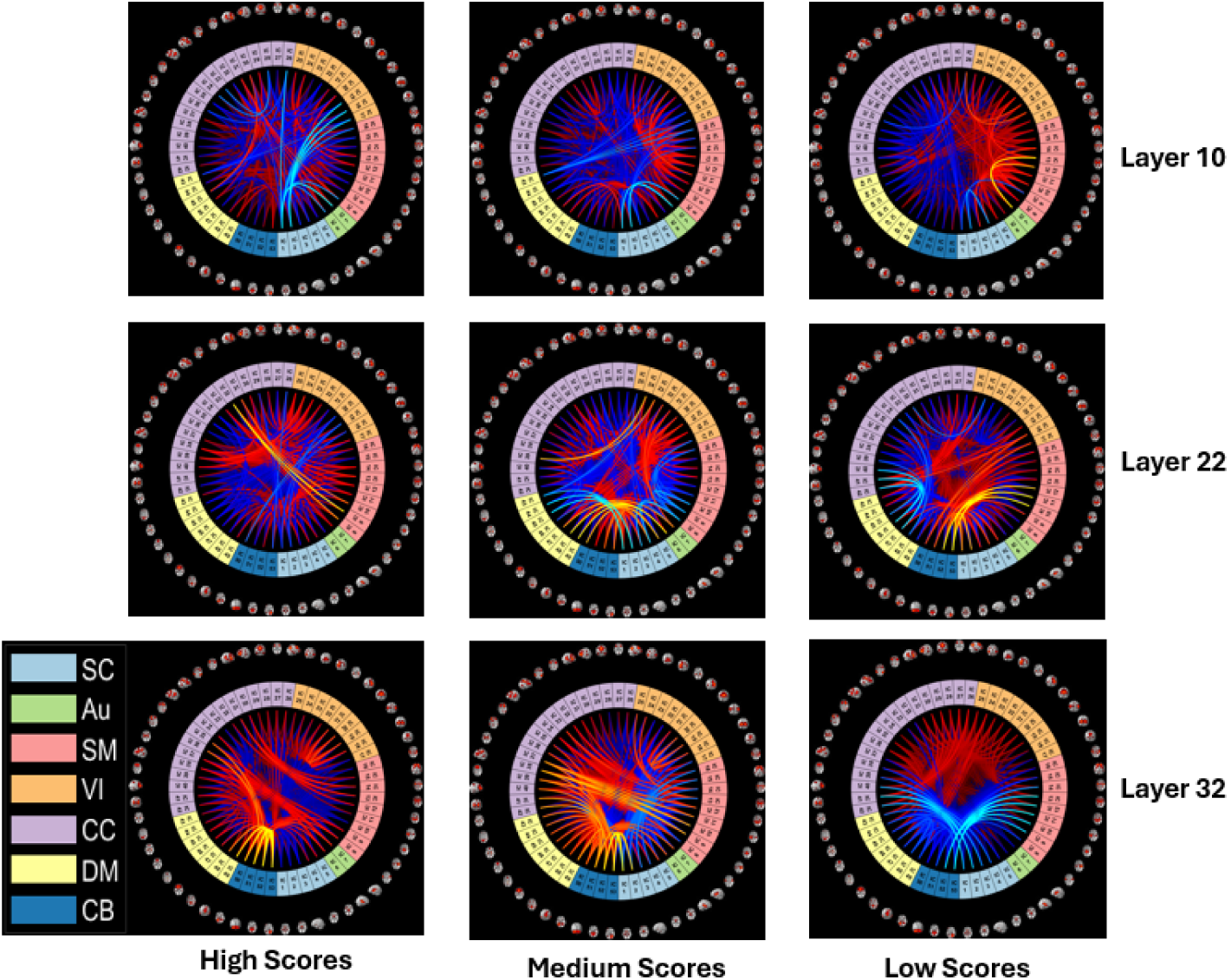
Connectograms of Two-sample t-tests for Layer 10, Layer 22, Layer 32 at different cognitive levels.

The figure displays a set of data that provides valuable insights into the differences between two groups of individuals. The mean values of the SZ group are significantly stronger when compared to the HC group in terms of the connections between the SC network and other networks, such as AUD, SM, VS, CC, and DM. These strong connections are evident at different cognitive levels, indicating a consistent pattern across different stages of cognitive processing.

Further analysis revealed that at high and medium cognitive levels, the HC group also showed evidence of more robust connections between the DM, CC, and CB networks. However, at the low cognitive level, the data shows that the mean values of the SZ group are significantly stronger compared to the HC group in these connections. This finding suggests that individuals with SZ may have a more heightened sensitivity to certain cognitive tasks, particularly those involving the DM, CC, and CB networks, which may contribute to the differences observed between the two groups. In Figure 14, you can take a look at the connectograms that highlight the most significant connections that exist in two separate networks - SC and CB.

Our study aimed to examine whether there exist any significant differences in means between two groups of individuals diagnosed with schizophrenia at different cognitive levels. To achieve this objective, we conducted two-sample t-tests on three different layers, which were 10, 22, and 32, as illustrated in Figure 15. The results displayed in this figure show that the mean values of the group diagnosed with SZ at the high cognitive level are significantly lower in the connections between SC, AUD, VM, and VS compared to the group of SZ at the low cognitive level. On the other hand, our analysis revealed that the SZ group at the low cognitive level has a higher mean than the SZ group at the high cognitive level in the VS and DM within networks.

**Figure 15:**
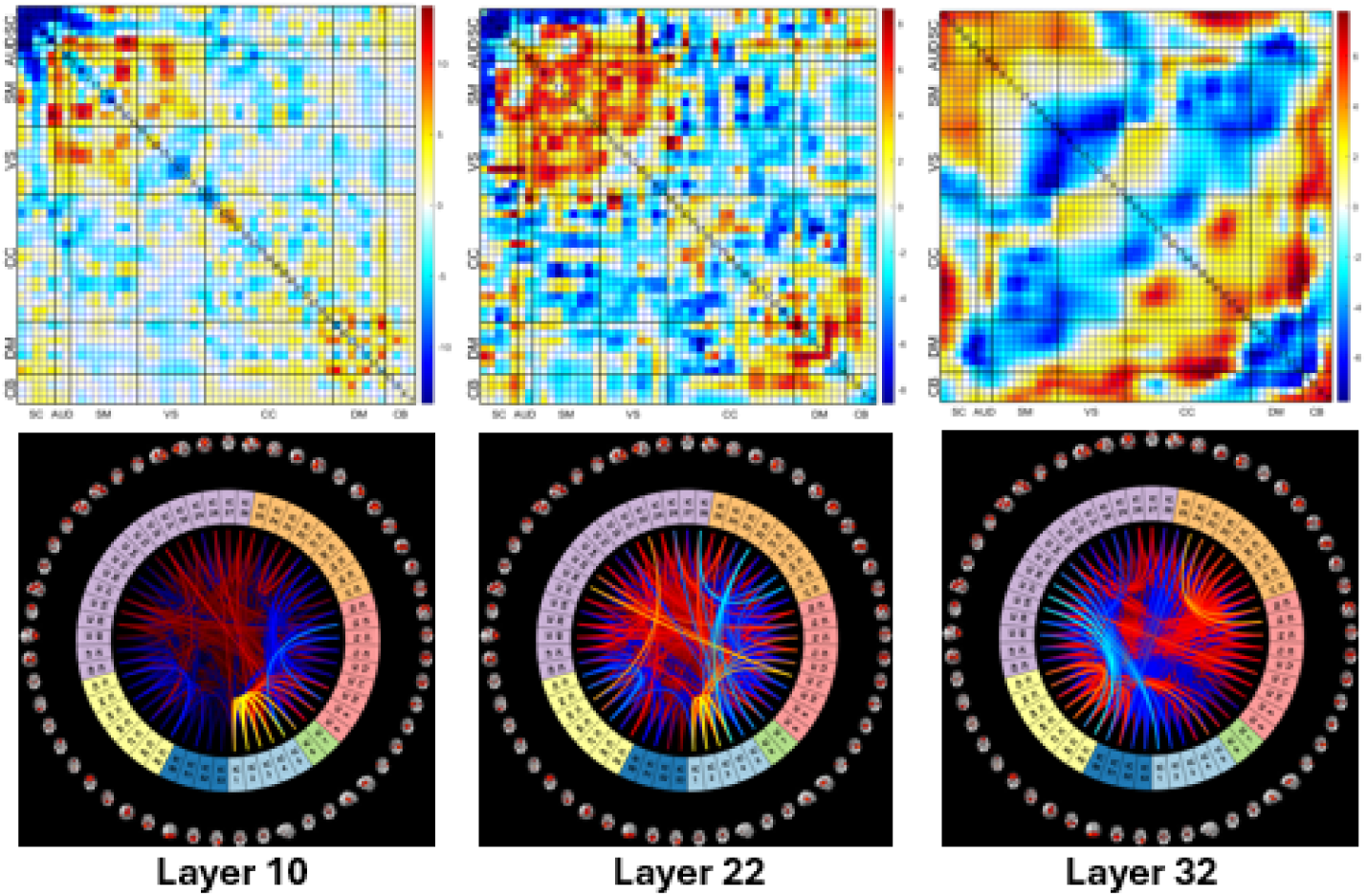
Differences in means between two groups of individuals diagnosed with schizophrenia at different cognitive levels.

### 7.9. Discussion

In recent years, there has been growing interest in utilizing feature selection techniques to better understand the abnormal brain functional connectivity that characterizes schizophrenia, using data obtained through functional magnetic resonance imaging (fMRI). Despite the significant progress made in this field, the underlying neurological mechanisms that contribute to the development of this disorder remain poorly understood. We have developed a deep-learning-based feature selection method that has demonstrated its ability to identify nonlinear and discriminative features in heatmaps with a high degree of accuracy. By leveraging the power of machine learning, our approach promises to shed new light on the complex neurobiological processes that underlie schizophrenia.

The paper we present outlines a new and innovative approach for extracting nonlinear heatmaps from functional network connectivity images. To do this, we utilize a deep convolutional neural network that effectively extracts these heatmaps from the input FNC data. By analyzing these heatmaps, we can derive remarkable nonlinear intrinsic connectivity networks that provide a robust framework for understanding the complex interactions between different regions of the brain. One of the key benefits of this approach is that it helps remove abundant information and noise from the heatmaps, leaving only the most important features related to identifying SZs. This means that these networks represent a significant improvement over previous approaches and offer a more accurate way of identifying SZs.

The heatmap generation process is crucial in identifying the most informative nonlinear functional connectivities that help distinguish between different groups. To achieve this, we have employed two-sample t-tests and ICA methods to analyze each heatmap from various convolution layers of the network. By doing so, we are able to extract the most relevant and significant nonlinear functional connectivities that contribute towards the classification of different groups. Overall, this new approach has the potential to greatly enhance our understanding of the brain and provide better insights into how different regions of the brain interact with each other.

Our study involved a rigorous evaluation of the classification performance of our proposed method as compared to the existing state-of-the-art classification methods. We used complex and challenging datasets to test the performance of our method, and the results were impressive. Our deep learning model was able to improve the classification results significantly, which demonstrated the effectiveness of our approach in identifying a small but crucial number of discriminative features to accurately classify different groups. Thus, our method was able to identify more discriminative features, which effectively helped in separating different classes. In comparison with the competing methods, we were able to showcase the superiority of our method through our results. In particular, our method has shown remarkable results in accurately identifying SZs and HCs. To be specific, we achieved an average accuracy of 92.8 % on the testing fMRI dataset, which is significantly higher than all other competing methods that had accuracies around 80 %. Moreover, our method’s ability to identify SZs and HCs at certain cognitive levels is a unique feature that cannot be achieved by other methods. This makes our method a promising tool for identifying and distinguishing between individuals with schizophrenia and healthy controls, which can have significant implications in the field of clinical psychology and neuroscience.

Through the process of identifying the most distinctive features from the heatmaps, we can delve into the dissimilarities in functional network connectivities between individuals who have been diagnosed with schizophrenia and those who have not (HC). After conducting our analysis, we have found that in comparison with the healthy control group, patients with SZ demonstrate notable differences in the following areas:

First, individuals diagnosed with schizophrenia demonstrate stronger connectivity between the sub-cortical network and other networks such as auditory, visual, sensorimotor, and cognitive control networks than the group of healthy individuals. This suggests that there may be an increased interaction between the sub-cortical network and other brain regions in individuals with SZ, potentially indicating a distinct pattern of brain functioning in this population.

Second, individuals diagnosed with schizophrenia exhibit a higher degree of connectivity between the default-mode network and other networks, such as visual and cognitive control networks, as compared to the group of healthy individuals. This increased connectivity indicates that the brain networks in individuals with schizophrenia are more tightly integrated, which could contribute to the symptoms associated with the disorder, such as hallucinations and delusions. The default-mode network is a set of brain regions that are active when the mind is at rest and not involved in any specific task, while the visual and cognitive control networks are responsible for visual processing and cognitive control, respectively. Third, the average values of the group diagnosed with SZ reveal a significant decrease in comparison to the group of HC in two types of brain connections - between-network connections and within-network connections - in the regions of sensorimotor, visual, and cognitive control. This means that individuals with SZ showcased lower connectivity levels in comparison to healthy individuals in the aforementioned regions of the brain.

Finally, upon conducting further analysis, it was discovered that at high and medium levels of cognitive function, the HC (healthy control) group displayed more pronounced connections between the default-mode, cognitive control, and cerebellum networks. These connections were found to be stronger in the HC group compared to the SZ group. However, at a lower level of cognitive function, the data showed that the mean values of the SZ group were significantly stronger in these connections when compared to the HC group. This finding suggests that individuals with SZ may have a heightened sensitivity to certain cognitive tasks, particularly those involving the default-mode, cognitive control, and cerebellum networks. These differences in neural connectivity may contribute to the observed variations between the two groups in cognitive abilities.

## 8. Conclusion

In this work, we have proposed a novel approach that harnesses the power of deep convolutional neural networks to extract nonlinear heatmaps from functional network connectivity images. Our approach is specifically designed to capture the complex nonlinear interactions that underlie cognitive operations and the changes observed in psychiatric conditions like schizophrenia. By doing so, we believe that this research can lead to a better understanding of the workings of the brain and offer crucial insights into the mechanisms that drive various psychological and psychiatric conditions.

Our method involves analyzing the heatmaps generated by the DCNN to derive nonlinear intrinsic connectivity networks from the corresponding input FNC data. These networks represent a significant improvement over previous approaches and provide a robust framework for understanding the complex interactions between different regions of the brain. To ensure optimal efficiency and effectiveness, our method incorporates two stages in the training process. In the initial stage, we train a deep convolutional neural network to create heatmaps from various convolution layers of the network. In the second stage, we use the ICA and t-tests-based feature selection methods to effectively analyze each heatmap from different convolution layers. This allows us to extract the most important nonlinear FNCs from the heatmaps that play a crucial role in distinguishing different groups. Overall, we believe that our approach can provide powerful insights into the workings of the brain and help researchers gain a deeper understanding of complex psychological and psychiatric conditions.

Our approach was focused on the use of nonlinear models to analyze FNC matrices. We find, even in this case, substantial benefits, including an improvement in accuracy and also a rich set of visualizations that highlight different aspects of the patient versus control differences. There are likely additional performance improvements to be gained by modeling nonlinear interactions prior to the computation of the FNC, which we hope to explore in future work. In addition, we are planning to enhance the effectiveness of our deep-learning network by incorporating self-attention transformers. This specialized transformer architecture is capable of capturing nonlinear connectivity from dynamic functional network connectivity with greater accuracy and precision. By enabling the model to recognize and analyze relationships between distant elements in a sequence, self-attention transformers can effectively comprehend complex patterns and dependencies that would be otherwise difficult to detect. This allows for the extraction of the most significant features from windowed time courses of different brain networks (components), which are estimated via spatial independent component analysis (sICA). We believe that this development will significantly improve the performance of our deep-learning network and enable us to achieve more accurate results in our research.

## Notes

### Competing Interest Statement

The authors have declared no competing interest.

